# Structure-activity Relationship for Diarylpyrazoles as Inhibitors of the Fungal Kinase Yck2

**DOI:** 10.1101/2025.07.12.664496

**Authors:** David J. Shirley, Bonnie Yiu, Ikeer Mancera-Ortiz, Peter J. Stogios, Zhongle Liu, Nicole Robbins, Luke Whitesell, Leah E. Cowen, David H. Drewry, Timothy M. Willson

## Abstract

Candida albicans is a growing global health threat, causing 1.5 million invasive infections and 1 million deaths annually. Yeast casein kinase 2 (Yck2) in *C. albicans* has emerged as an antifungal target of the kinase inhibitor LY364947 (**LY**). Herein, we report Yck2 structure-activity relationships for 3,4- and 3,4,5-substituted pyrazole analogs of **LY**. X-ray crystallography and in vitro profiling revealed the importance of the hinge-binding heterocycle for Yck2 inhibition and fungal kinome selectivity. A hydrogen-bond network between the inhibitor, a bound water molecule, and catalytic residues within the ATP pocket was identified as a key determinant of selectivity over other fungal and human kinases. Phenol analog **11** showed remarkable selectivity for Yck2 and Yck22 over all other *C. albicans* protein kinases. Several of the **LY** analogs, including **11**, demonstrated improved antifungal activity. These findings provide a framework for translating human kinase inhibitors into highly selective antifungal Yck2 inhibitors.

**Graphical Abstract:** 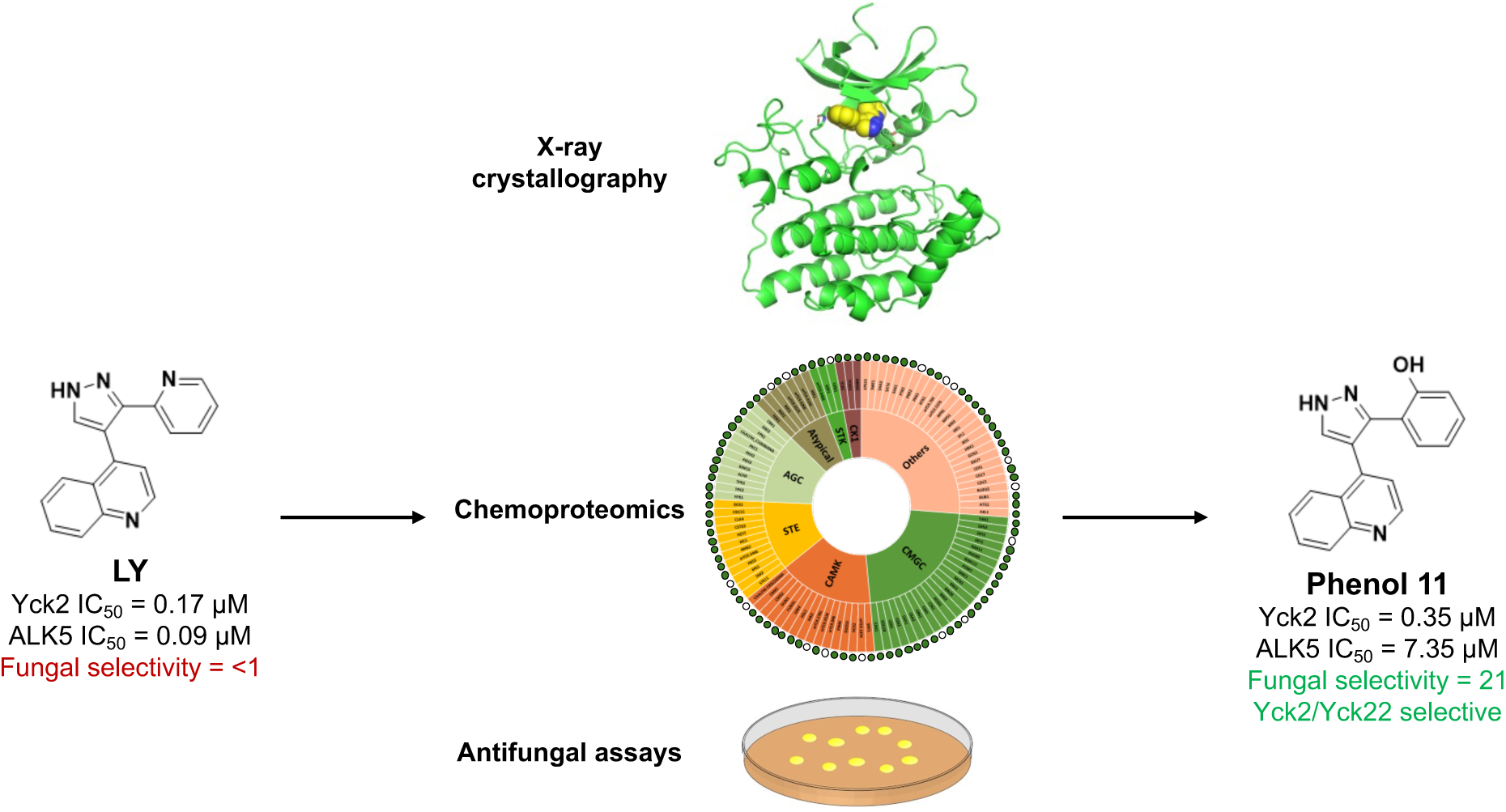

---

Yeast Casein Kinase 2 (Yck2) is a stress response protein kinase that helps regulate key features of *C. albicans* pathogenicity, including fungal cell wall integrity, biofilm formation, and actin-related processes during morphogenesis.^1,2^ Yck2 was identified in *C. albicans* as a primary target responsible for the antifungal activity of the kinase inhibitors GW461484A (**GW**), YK-1-02 (**YK**), and MN-1-157 (**MN**). **YK** and **MN** also demonstrated antifungal activity in a mouse model of systemic candidiasis, reducing fungal burden as single agents and in combination with the echinocandin caspofungin.^3,4^ The pyrazolo[1,5-*a*]pyridine chemotype found in **GW** and **YK** was originally developed for targeting of human MAPK14 (a.k.a. p38α) but was successfully repurposed for development of fungal-selective Yck2 inhibitors (Figure 1).^5,6^ When profiled across the *C. albicans* kinome, **GW** and **YK** demonstrated remarkable selectivity for Yck2 and its paralogs Yck22 and Hrr25,^7^ with only two additional off-target kinases: Hog1 (the fungal MAPK14) and Pom1 (an ortholog of the dual-specificity tyrosine-regulated kinase/DYRK).^8,9^ Yck2, Yck22, and Hrr25 are the fungal orthologs of the eukaryotic casein kinase 1 (CK1) family.^10,11^ A screen of known human CK1 inhibitors with selectivity over MAPK14 to identify new fungal Yck2 inhibitors led to the diarylpyrazole LY364947 (**LY**), which was originally synthesized as a human ALK5 inhibitor.^7,12^ Another human CK1 inhibitor **MU1742** with high selectivity over MAPK14 that combines features of both **GW**/**YK** and **LY** chemotypes was recently reported.^13^ However, **MU1742** displayed surprisingly poor activity against Yck2 despite its potent inhibition of human CK1, demonstrating the challenge of translating SAR from human to fungal kinases.^7^ Our success in repurposing inhibitors of human CK1, MAPK14, and ALK5 for use as fungal Yck2 inhibitors has further highlighted the need to fully investigate the chemogenomic relationship between the fungal and human kinome. In this study we set out to define structure-activity relationships for inhibition of *C. albicans* Yck2 by the diarylpyrazole chemotype found in **LY** and to identify the similarities and differences between its fungal and human kinome profiles.

**Figure 1.**
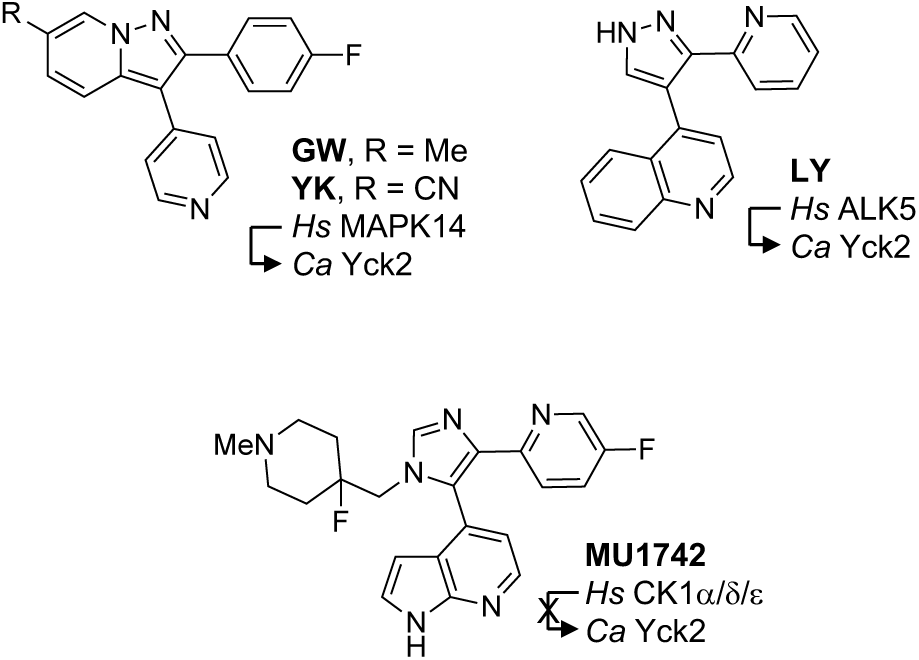
Repurposed human kinase inhibitors. *Hs*, *Homo sapiens*. *Ca*, *C. albicans*. Alignment of the amino acid sequences of *Ca*Yck2, *Hs*CK1α, *Ca*Hog1, *Hs*MAPK14, and *Hs*ALK5 showed that while identity across the kinase *N*-lobe was low, many residues within key motifs of the ATP-binding pocket were highly conserved (Table S1).^14–18^ To understand the full extent of the differences in the pocket residues between the human and fungal kinases the X-ray crystal structure of **LY** in complex with Yck2 was solved. Crystallization was performed using the sitting drop method with **LY** soaked into *apo*-Yck2 crystals over 24 h. X-ray diffraction collection on a RigakuMicroMax-003 and structure refinement using Phenix^19,20^ resulted in a 1.7 Å resolution structure (PDB: 9NZK; Figure 2a and 2b, Table S2).

Comparison of the structure of **LY** in complex with Yck2 (Figure 2b) with its previously determined structure^8^ bound to ALK5 (Figure 2c) showed that the binding orientation was nearly identical. Common interactions of the inhibitor within the ATP-binding pocket were seen in both structures. The quinoline nitrogen formed an H-bond to a backbone amide of the hinge-binding region (L120 in Yck2, H283 in ALK5) (Figure 2b and 2c). The hinge region, which has the lowest sequence similarity across protein kinases, appeared to have a significant impact on human kinase selectivity, specifically towards inhibitors with MAPK14 activity: original reports of **LY**^12^ and **MU1742**^13^ showed modification of their hinge-binding pharmacophore improved on-target (ALK5 and CK1α) potency and selectivity over MAPK14. Although both Yck2 and MAPK14 engage a leucine in hydrogen-bonding *via* its amide backbone, none of the neighboring amino acids are identical, resulting in different electronic environments within the binding pocket. We used these results to design **LY** analogs featuring a range of hinge-binding heterocycles and to investigate their effect on fungal kinase selectivity.

**Figure 2.**
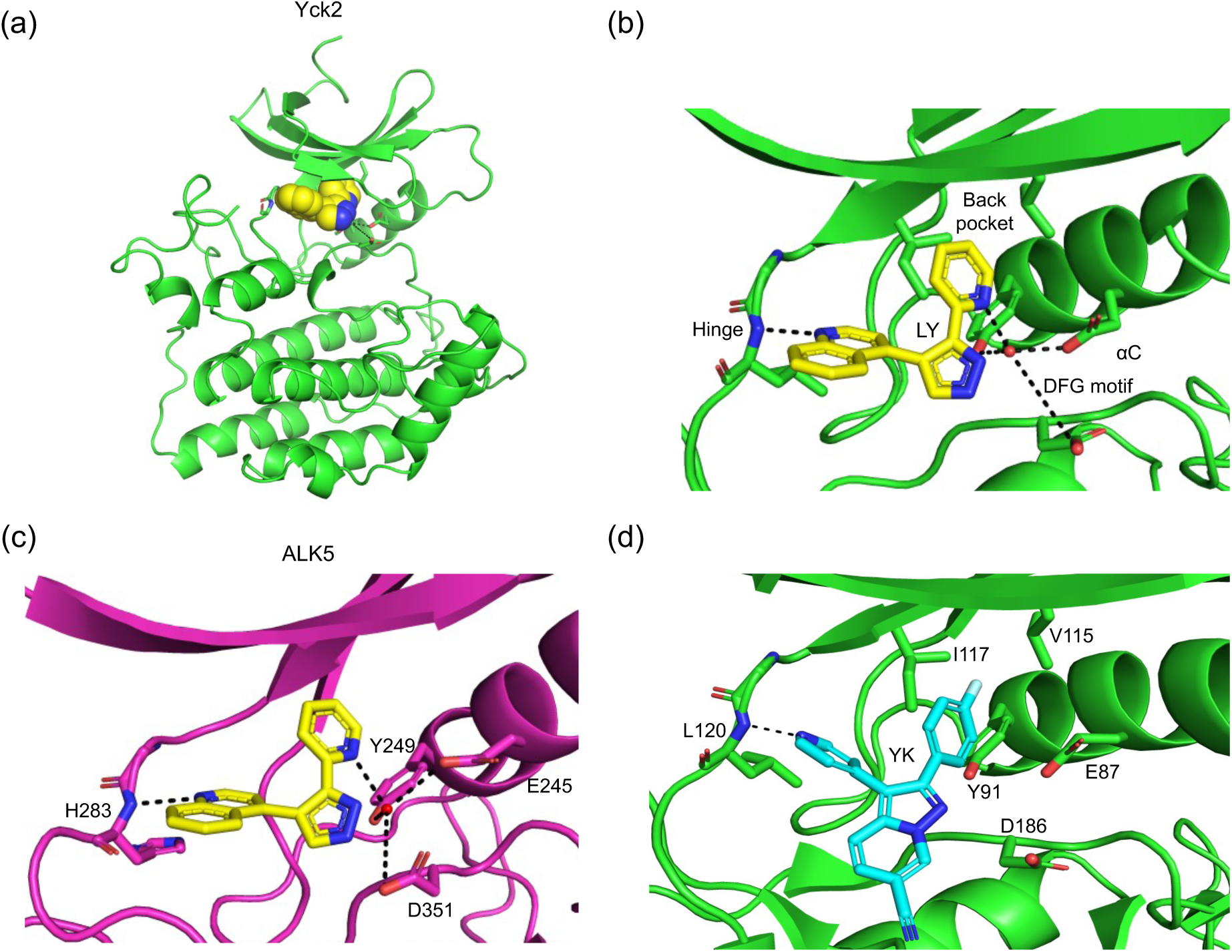
Crystal structures of *C. albicans* Yck2 and human ALK5 in complex with **LY** and **YK**. (a) View of Yck2 (green ribbon) bound to **LY** (yellow spheres) from PDB: 9NZK. (b) Key motifs in the Yck2 ATP-binding pocket of the structure with **LY** (yellow sticks). (c) ATP-binding pocket of ALK5 (magenta ribbon) bound to **LY** (yellow sticks) from PDB:1PY5. (d) ATP-binding pocket of Yck2 (green ribbon) bound to **YK** (cyan sticks) from PDB: 9EDX. H-bonds are portrayed as black dashes, with hinge-binding interactions (b, c, and d) and water-bonding networks indicated (b and c).

The Yck2-**LY** structure (Figure 2a and 2b) showed that the nitrogen of the 3-pyridyl group was engaged in a hydrogen bond to a water molecule that bridged to a glutamate residue in the conserved αC-helix (E87 in Yck2, E285 in ALK5), a neighboring tyrosine (Y91 in Yck2, Y249 in ALK5), and an aspartate residue D186 in the DFG motif (D351 in DLG motif for ALK5). One subtle difference in the structures was the distance between the bound water molecule and the aspartate of the DFG-motif (5.9 Å in Yck2 vs. 4.9 Å in ALK5), which suggested a weaker interaction in the fungal kinase. This bound water molecule may be important for maintaining Yck2 potency and MAPK14 selectivity, since removal of the pyridine nitrogen caused a substantial drop in potency for ALK5 and maintained MAPK14 selectivity in the **LY** series.^8^ Notably no bound water molecule was observed in the Yck2-**YK** co-crystal structure (Figure 2d), which may account for its off-target inhibition of Hog1, the fungal MAPK14.^7^

The 3-pyridyl group of **LY** occupied space near the gatekeeper residue and the hydrophobic back pocket (Figure 2b). The back pocket is relatively conserved across protein kinases, both in sequence and function, but the gatekeeper residue is unique for every kinase in our sequence alignment (Table S1). Yck2 and CK1α are the most similar across the kinase gatekeeper residues with an isoleucine (I117) and methionine (M90), respectively, while the other kinases contain residues with polar sidechains (S280 in ALK5, T106 in MAPK14, N100 in Hog1; Table S1). Although the gatekeeper and back pocket of Yck2 have lipophilic residues (I117 and V115) that could engage in hydrophobic interactions, there is only limited space to extend **LY** into this pocket (Figure 3a and b). The back pocket opening is ∼6 Å deep and 3.7 Å at its widest, restricting distal extension and aromatic substitution to small chemical motifs. Both fungal Yck2-selective (**YK**) and mammalian CK1-selective (**MU1742**) inhibitors contain *m*-fluoroaryl groups, highlighting the importance of small, lipophilic substituents to engage the back pocket in these kinases.

**Figure 3.**
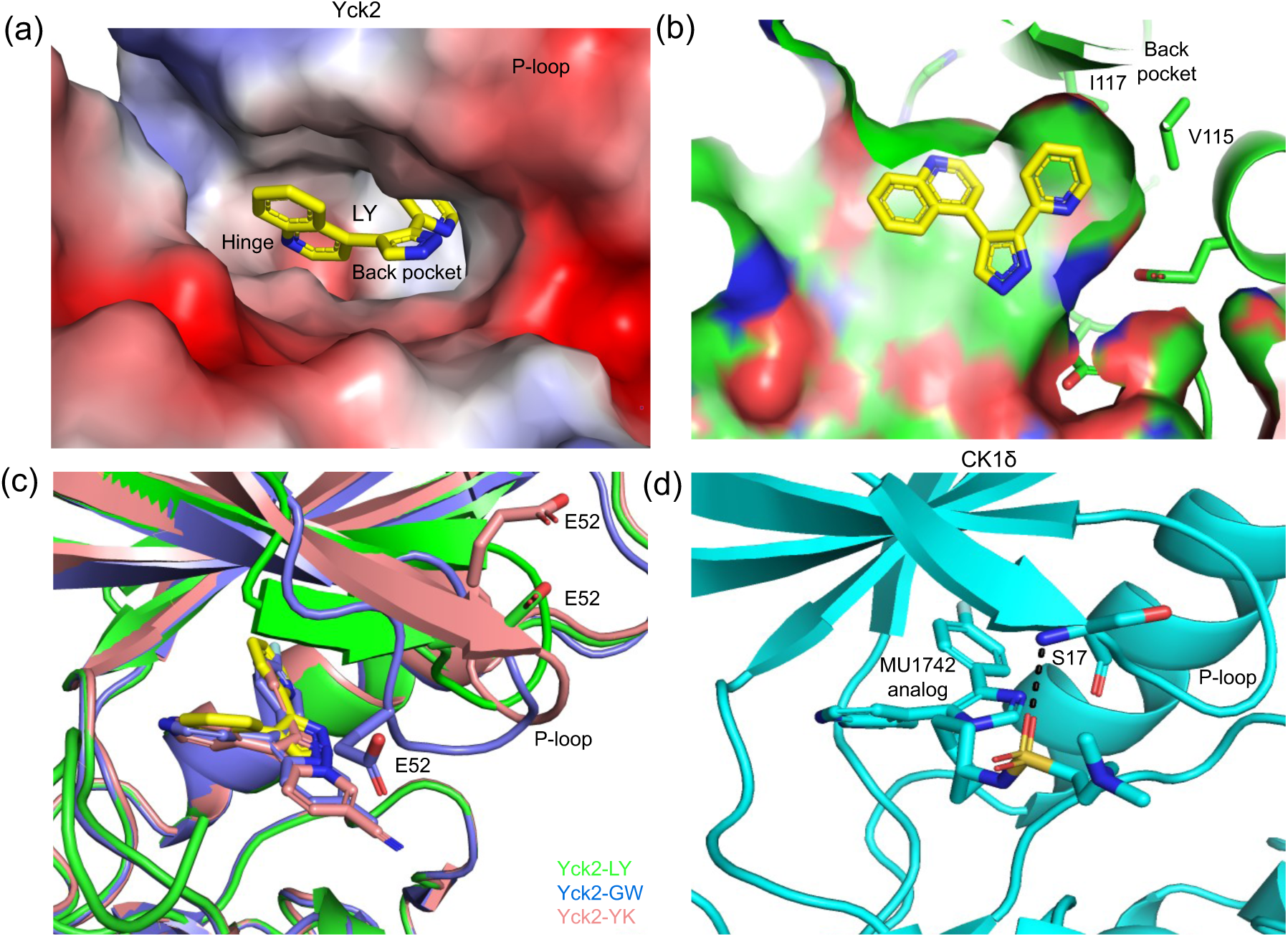
Analysis of key motifs within the ATP-binding pocket. (a) Topological analysis of the Yck2 back-pocket. Inward view of Yck2 electrostatic surfaces (red: acidic, blue: basic) bound to **LY** (yellow sticks) from PDB: 9NZK. (b) Cross-sectional view of the Yck2 back-pocket (green surface) depicting key residues, and the orientation of the C-3 pyridine within the back pocket. (c) Multiple conformations of the glycine-rich P-loop. Overlay of Yck2 bound **LY** (green ribbon, PDB: 9NZK), **GW** (blue ribbon, PDB: 6U6A), and **YK** (peach ribbon, PDB: 9EDX) with different positions of P-loop E52 shown. (d) CK1δ bound to **MU1742** (cyan ribbon, PDB:7QRB) with the H-bond between the inhibitor and the P-loop S17 shown.

Sequence alignment of Yck2 with CK1α, MAPK14, ALK5, and Hog1 showed substantial overlap in key ATP-binding motifs (Table S1). However, the flexible glycine-rich P-loop exhibited poor conservation of amino acids. We attempted to engage the glutamate residue of the Yck2 P-loop (E52) with basic **YK** and **GW** derivatives, but the resulting X-ray structures showed that the flexible loop adopted multiple conformations confounding our initial inhibitor design (Figure 3c). Notably, an **MU1742** analog with a C-5 sulfonamide was shown to interact with the CK1δ P-loop through a hydrogen bond network with a water molecule and S17 (Figure 3d).^13^ This interaction was responsible for improved human CK1-family kinase selectivity over MAPK14. Interestingly, **MU1742** was a poor inhibitor of fungal Yck2 for reasons that were not obvious from analysis of the Yck2-**LY** structure.^4^ In the **LY** chemotype, addition of a substituent at N-1 would mimic the *N*-methylpiperidine substituent on **MU1742** (Figure 4).

**Figure 4.**
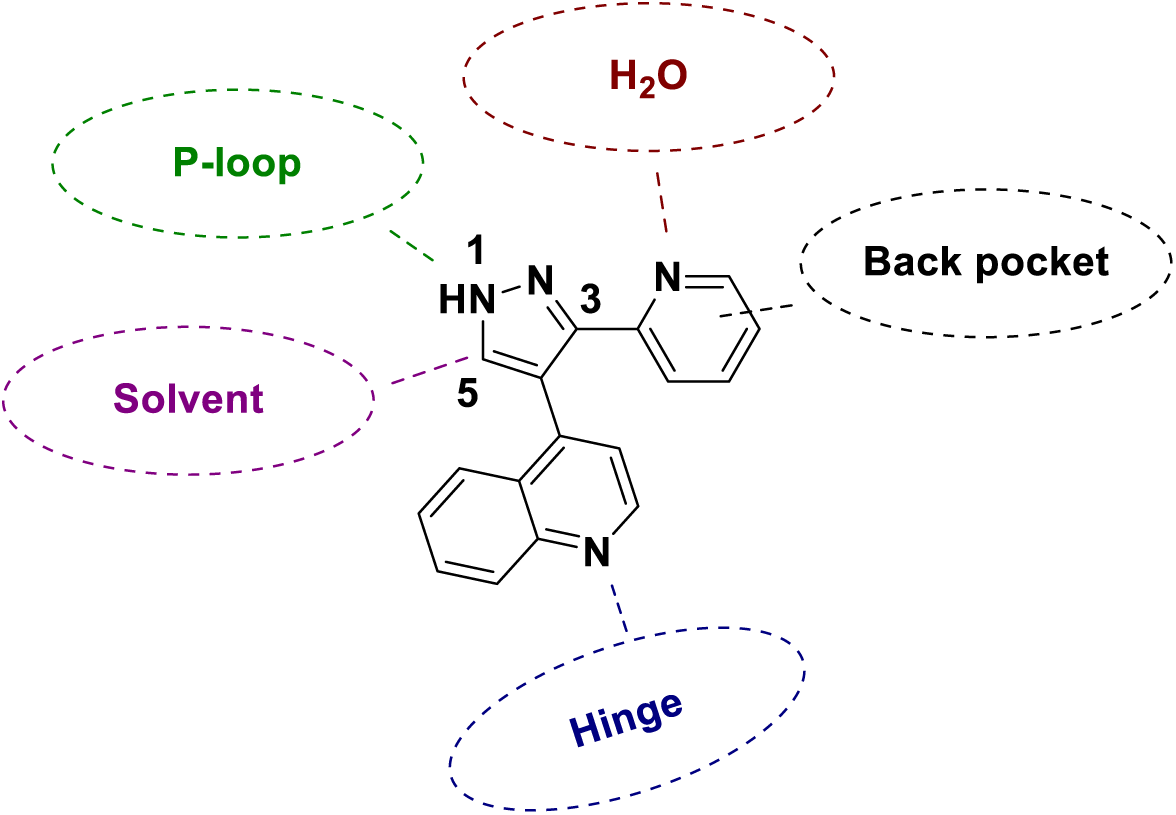
Design of **LY** analogs from analysis of X-ray structures. Hinge binding groups were selected to improve fungal Yck2 potency and selectivity. Substitutions at the C-5/N-1 position targeted the Yck2 P-loop. Modifications of the C-3 motif were designed to both probe the significance of the H-bond to the bound water molecule and explore the Yck2 back pocket.

New analogs of **LY** were synthesized using modifications of the reported methods (Scheme 1).^12^ Methyl quinoline-like scaffolds underwent 1,2-addition into picolinates, followed by the addition of hydrazine or hydrazide and heating to yield cyclized 3,4- or 3,4,5-substituted pyrazoles. Compounds with N-1 substitution (**1** and **2**) and C-5 substitution (**3** and **4**) were prepared. For modification of the pyrazole core, pyrrole (**5**) and furan (**6**) analogs were synthesized using enolate chemistry, ozonolysis, and subsequent cyclization. A handful of analogs were synthesized with alterations to the hinge binder, possessing either a pyridine (**7**) or pyrimidine (**8**) substituent. Alteration of the H-bond acceptor in the 3-aryl group was investigated through removal of the pyridine nitrogen (**9**), methylation (**10**), or replacement by a phenol or furan (**11** and **12**).

**Scheme 1.**
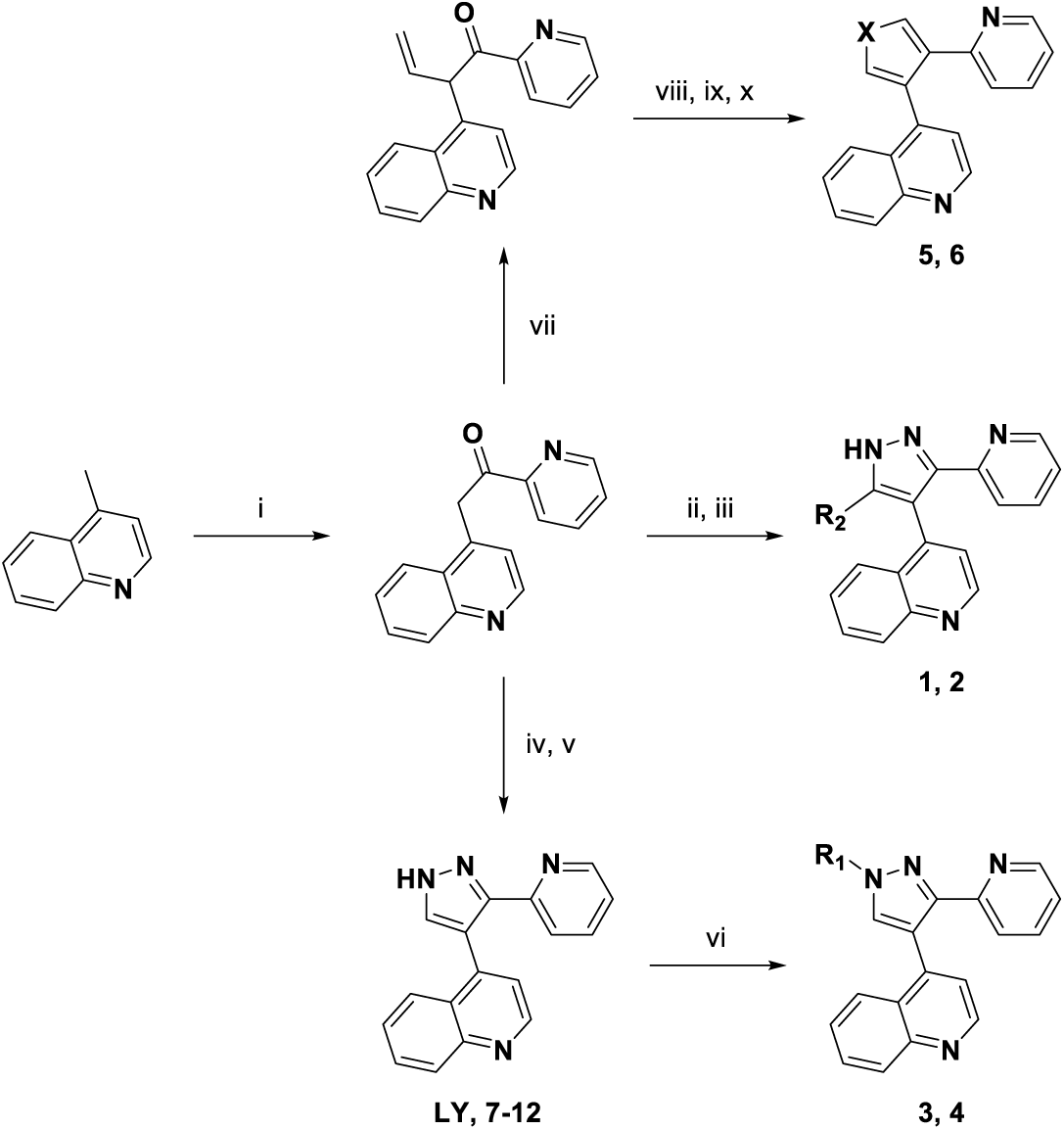
(i) methyl picolinate, LiHMDS, THF, 0 °C to rt, 18 h. (ii) H2NNH2, HCl, 40 °C, 3 h. (iii) DMF, microwave at 200 °C, 8 h. (iv) DMFDMA, THF, 60 °C, 18 h. (v) H2NNH2•H2O, EtOH, rt, overnight. (vi) alkyl halide, NaH, DMF, rt, 1 h. (vii) allyl bromide, LiHMDS, DMSO/THF, 0 °C, 3 h. (vii) O3, 10% MeOH/DCM, -78 °C, 2 min. (ix) X = NH, NH4CH3CO2, glacial acetic acid, microwave at 120 °C, 2 h. (x) X = O, conc. HCl, glacial acetic acid, microwave at 120 °C, 2 h.

Kinase enzyme inhibition was measured using ADP-Glo assays employing purified recombinant Yck2 kinase domain or full length CK1α and MAPK14 proteins (Tables 1–3). A series of analogs (**1**–**4**) with C-5 or N-1 substitution explored the role of the P-loop in determining fungal selectivity (Table 1). Although **MU1742** has a very specific *N-*methylpiperidine group, a simple C-5 benzyl substituted analog (e.g. **MU1442**) was reported to demonstrate CK1α IC50 <20 nM with 50-fold selectivity over MAPK14.^13^ In contrast, in the **LY** series, we found that C-5 benzyl (**1**) and *p*-fluorobenzyl (**2**) analogs had a 28- to 41-fold drop in potency for Yck2 inhibition as well as 10-fold or greater potency reduction for CK1α and MAPK14 inhibition. This reduction in potency from C-5 substitution may result from free rotation of the benzylic carbon, which is not tolerated in Yck2. N-1 substitution in analogs **3** (Me) and **4** (PhCH2) had a 3- to 6-fold drop in Yck2 potency, with a 2-fold improvement and 3-fold reduction, respectively for MAPK14 selectivity. The N-1 substituent faces the P-loop and the solvent-exposed region of the ATP binding pocket (Figure 4). The improved selectivity of analog **4** over MAPK14 as well as CK1α suggests that N-1 substitution may result in a favorable interaction with the flexible P-loop.

**Table 1.**

| Compound | X | Y | R<sup>1</sup> | R<sup>2</sup> | Yck2<br>IC<sub>50</sub> (μM)<sup>a</sup> | CK1α<br>IC<sub>50</sub> (μM)<sup>a</sup> | Selectivity<br>(CK1α/Yck2) | MAPK14<br>IC<sub>50</sub> (μM)<sup>a</sup> |
| --- | --- | --- | --- | --- | --- | --- | --- | --- |
| **LY** | N | N | H | H | 0.17 | 1.0 | 6 | 1.5 |
| **1** | N | N | H | PhCH<sub>2</sub> | 4.7 | 3.5 | 1 | 5.2 |
| **2** | N | N | H | 4-F-Ph | 7.0 | 11 | 2 | 3.5 |
| **3** | N | N | Me | H | 0.55 | 1.5 | 3 | 9.4 |
| **4** | N | N | PhCH<sub>2</sub> | H | 1.0 | 5.9 | 6 | 3.0 |
| **5** | C | NH | H | H | 4.9 | >30 | >6 | >30 |
| **6** | C | O | H | H | 6.4 | >30 | >5 | >30 |

<sup>a</sup>All data n = 1 in triplicate ±15%

**Table 2.**

| Compound | X | Yck2<br>IC<sub>50</sub> (μM)<sup>a</sup> | CK1α<br>IC<sub>50</sub> (μM)<sup>a</sup> | Selectivity<br>(CK1α/Yck2) | MAPK14<br>IC<sub>50</sub> (μM)<sup>a</sup> |
| --- | --- | --- | --- | --- | --- |
| **7** | C | 0.52 | 2.9 | 6 | 4.5 |
| **8** | N | 1.6 | >30 | >20 | 1.5 |

<sup>a</sup>All data n = 1 in triplicate ±15%

**Table 3.**

| Compound | R | Yck2<br>IC<sub>50</sub> (μM)<sup>a</sup> | CK1α<br>IC<sub>50</sub> (μM)<sup>a</sup> | Selectivity<br>(CK1α/Yck2) | MAPK14<br>IC<sub>50</sub> (μM)<sup>a</sup> |
| --- | --- | --- | --- | --- | --- |
| **9** | 4-F-Ph | 0.59 | 4.3 | 7 | 0.54 |
| **10** | 4-Me-2-pyridyl | 4.7 | >15 | >3 | 17.5 |
| **11** | 2-OH-Ph | 0.35 | 3.1 | 10 | 0.73 |
| **12** | 2-furyl | 0.83 | 3.4 | 5 | 1.8 |

<sup>a</sup>All data n = 1 in triplicate ±15%

Two compounds were designed to explore the role of the pyrazole core in Yck2 inhibition. Compared to **LY**, the pyrrole (**5**) and furan (**6**) had a 29- and 37-fold weaker Yck2 inhibition, respectively (Table 1). This reduction in Yck2 inhibition was unexpected, since **5** and **6** were designed as bioisosteric replacements of the pyrazole. Their heteroatoms were expected to be solvent exposed with only one hydrophobic residue (I54) within 3 Å. Changing the heterocyclic core may result in a modified orientation of the quinoline and pyridine substituents that is detrimental to Yck2 inhibition.

Two analogs explored the contribution of the hinge-binding group to fungal kinase inhibition potency and selectivity (Table 2). SAR conducted on **LY** by Eli Lilly showed weaker H-bond acceptors maintained good potency for ALK5, while stronger H-bond acceptors led to increased MAPK14 potency.^12^ Notably, the results seen in Eli Lilly’s report are mirrored here. Pyridine (**7**) and pyrimidine (**8**) analogs of **LY** demonstrated a small reduction in potency (3- to 10-fold). Despite the reduction in Yck2 potency, **7** maintains the same degree of selectivity for Yck2 over CK1α and MAPK14 compared to **LY**. The stronger hydrogen-bond acceptor **8** displayed a 5-fold improvement in fungal selectivity over CK1α. Unfortunately, neither analog had improved selectivity over MAPK14. Additional investigation of alternative hinge-binding groups may be necessary to enhance Yck2 potency and MAPK14 selectivity.

The role of the variable gatekeeper residue and its impact on the size of the kinase back pocket was explored using several analogs with modified substituents at C-3 (Table 3). Crystal structures show that the Yck2, and CK1α, back pocket is very shallow while the MAPK14 back pocket is slightly larger (∼7 Å deep, 4 Å wide). Small lipophilic substitutions in analogs **9** (4-F) and **10** (4-Me) were designed to explore these differences. Methylation of the pyridine (**10**) caused a 27-fold drop in potency indicating a steric clash with the smaller Yck2 back pocket. The 4-fluorophenyl analog **9**, designed to engage in hydrophobic interactions with the gatekeeper residue (I117) and neighboring leucine (L115) (Figures 2b, 3b and 4), maintained sub-micromolar potency for Yck2 even with the absence of the key 3-pyridyl nitrogen. Both CK1α and ALK5 possess a tyrosine on the αC-helix engaged in a hydrogen-bonding network with a water molecule and the pyridine in **LY** (Figure 2b and c). The 3-phenol (**11**) and furan (**12**) derivatives demonstrated comparable Yck2 potency to **LY** but with poor selectivity over MAPK14. Notably, MAPK14 does not possess a tyrosine in its αC-helix, so further investigation is required to identify interactions responsible for maintaining the potency of **11** and **12** on this kinase.

**LY** and analogs **3**, **9**, **11**, and **12** were tested for inhibition of ALK5 to further assess their kinase selectivity (Table 4). **LY** was ∼2-fold more potent for inhibition of ALK5 compared to Yck2. However, analogs **3**, **9**, **11**, and **12** showed improved fungal Yck2 selectivity when compared to the parent analog. While the methylated pyrazole core (**3**) and C-3 furan (**12**) had equivalent Yck2 and ALK5 inhibition, the C-3 fluorophenyl (**9**) analogs showed >10-fold improvement in fungal selectivity over ALK5 compared to **LY.** However, most remarkable was the C-3 phenol (**11**) analog, which showed >40-fold improvement in selectivity over ALK5. This dramatic improvement may be due to a stronger H-bond network formed with the phenolic hydroxyl group compared to the corresponding 3-pyridyl group in the Yck2-**LY** and ALK5-**LY** structures (Figure 2b and c).

**Table 4.**

| Compound | Yck2<br>IC<sub>50</sub> (μM) | ALK5<br>IC<sub>50</sub> (μM)<sup>a</sup> | Selectivity<br>(ALK5/Yck2) |
| --- | --- | --- | --- |
| **LY** | 0.17 | 0.09 | <1 |
| **3** | 0.55 | 0.75 | 1 |
| **9** | 0.59 | 3.7 | 6 |
| **11** | 0.35 | 7.4 | 21 |
| **12** | 0.83 | 1.1 | 1 |

<sup>a</sup>All data n = 1 in triplicate ±15%

To assess the kinome-wide fungal selectivity of the **LY** analogs, chemoproteomic analysis was conducted with **11** and **12** in *C. albicans* lysate using multiplexed inhibitor beads coupled with mass spectrometry (MIB/MS) analysis (Figure 5).^7,21,22^ We previously described application of this chemoproteomic method to capture the *C. albicans* kinome and measure the fungal selectivity of **YK** and **LY**.^7^ Phenol **11** was selected for kinome profiling based on its Yck2 potency being the best among all analogs synthesized (IC50 = 0.35 μM), along with its improved CK1α (10-fold) and ALK5 (21-fold) selectivity. Furan **12** was selected for MIB/MS analysis because it had good Yck2 potency and possessed an alternative functionality at the C-3 position. The MIBS matrix captured 86 *C. albicans* protein kinases, representing 83% of the fungal protein kinome (Figure S1).^23^ MIB/MS competition experiments were performed with **11** and **12**. Hits were defined as kinases exhibiting a dose dependent Log2 fold change ≤ -1.0.

**Figure 5.**
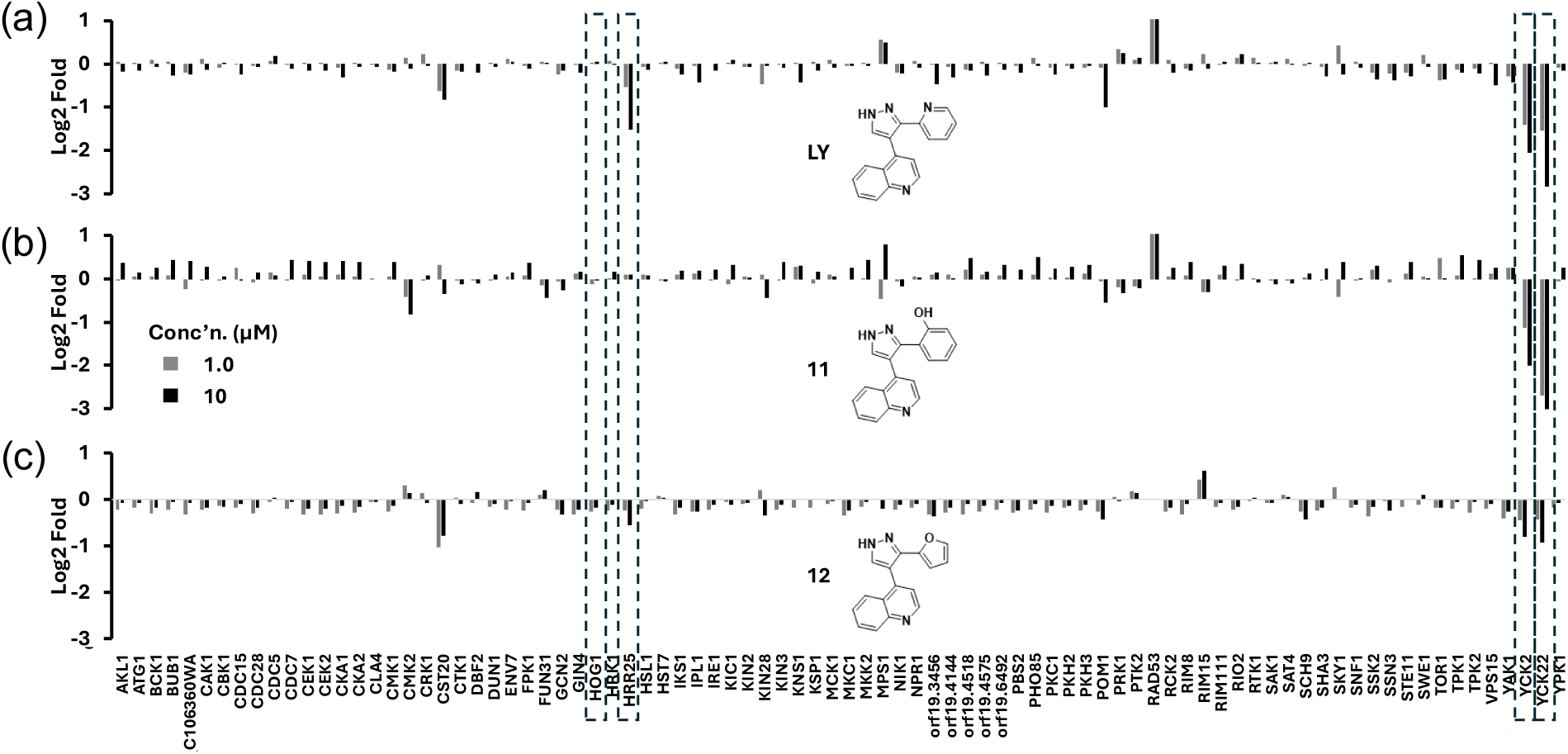
Fold-change of fungal protein kinase targets captured in MIB/MS competition assays for each compound relative to DMSO (a) Competition with **LY**. (b) Competition with phenol **11**. (c) Competition with furan **12**. Hits are defined as kinases exhibiting a dose dependent Log2 fold change ≤ -1.00. n = 1. Hog1, Hrr25, Yck2, and Yck22 are highlighted by the hatched boxes.

Phenol **11** displayed competition with Yck2 comparable to **LY**, however, there were noticeable differences in its kinase selectivity. Phenol **11** exhibited a 2-fold increase in potency against Yck2 paralog Yck22 but also demonstrated selectivity over paralog Hrr25. This result is the first report of a Yck2/Yck22 inhibitor with selectivity over Hrr25. Phenol **11** was also selective over all other fungal protein kinases indicating that it is the most selective Yck2/Yck22 inhibitor reported to date. Compared to **LY** in the MIBS competition assay, furan **12** showed a 2-fold and 4-fold reduction in Yck2 and Yck22 competition, respectively. This result was anticipated based on the modest Yck2 biochemical activity of **12**. It is interesting that **LY**, **11**, and **12** did not compete for binding of Hog1, despite demonstrating modest potency for inhibition of human MAPK14. Based on primary sequence analysis, Hog1 and MAPK14 share many residues at key binding motifs, with the greatest variation at the hinge and αC-helix (Table S1). These findings highlight distinct ligand-kinase binding differences between fungal and human kinases.

To investigate antifungal activity of the **LY** analogs, growth of wild-type *C. albicans* (CaSS1) was measured in the presence of increasing doses of **1**-**12**. These experiments were performed growing in RPMI medium at 37 °C under 5% CO2, conditions that mimic host conditions under which Yck2 is required for growth. At 100 μM, only **LY**, **2**, **9**, and **10** inhibited growth (Figure 6a). Additional assays were performed with a strain of *C. albicans* where one allele of *YCK2* was deleted and the other allele placed under transcription control of a strong, DOX-repressible promoter to validate the Yck2-dependence of the antifungal activity (*tetO-YCK2/yck2Δ*) (Figure 6b). Many of the **LY** analogs, including **LY**, **3**, **7**, and **9**, showed robust inhibition of fungal growth in the *tetO-YCK2/yck2*Δ strain when doxycycline was added to the culture medium, indicating that they were able to inhibit fungal growth when expression of *YCK2* was partially reduced. Notably, **1**, **2**, and **6** with Yck2 IC50 >4.0 µM remained inactive even in the *tetO-YCK2/yck2*Δ strain. To investigate the basis of the poor wild-type whole cell activity of the **LY** analogs, growth was measured using *C. albicans* strains modified to delete major drug efflux pumps (*cdr1Δ/Δ, cdr2Δ/Δ, mdr1Δ/Δ, flu1Δ/Δ)* or to alter fungal membrane permeability (*tetO-ERG6/erg6Δ)* (Figures 6c and 6d). Deletion of the efflux pumps (Figure 6c) had modest or no effect on the growth-inhibitory activity of compounds. However, eight of the twelve analogs had a >30-fold increase in the potency of antifungal activity when cell membrane permeability was increased by reducing expression of *ERG6*, a gene involved in the biosynthesis of ergosterol (Figure 6d).

**Figure 6.**
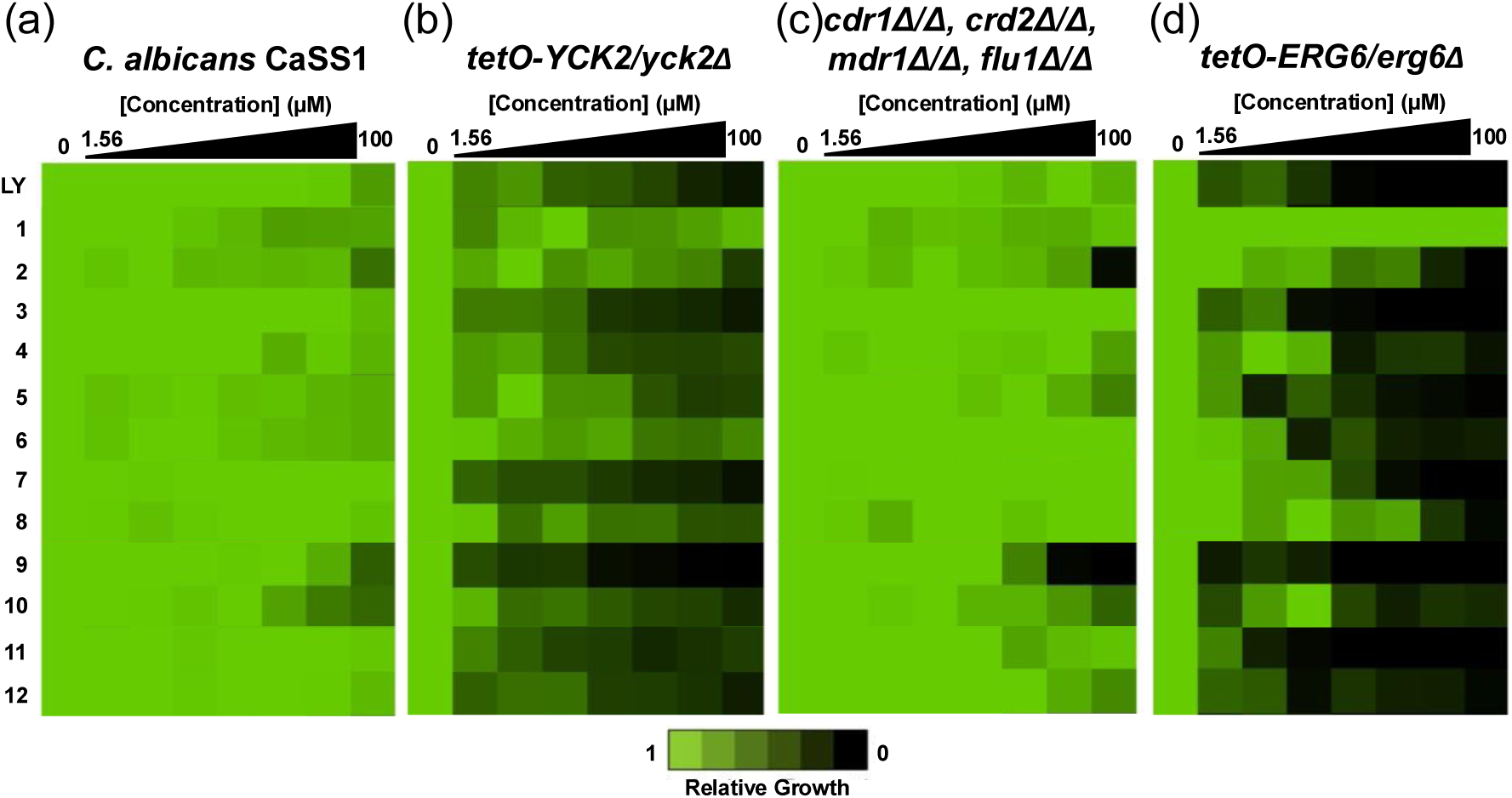
*C. albicans* antifungal activity of **LY** and analogs **1**–**12**. (a) Wild-type CaSS1. (b) *tetO-YCK2/yck2Δ*. (c) *cdr1Δ/Δ, cdr2Δ/Δ, mdr1Δ/Δ, flu1Δ/Δ*. (d) *tetO-ERG6/erg6Δ*. Growth was determined by optical density at 600 nm (OD600) with technical and biological duplicated normalized to compound-free controls (see color bar).

In summary, analogs of **LY**, designed through analysis of multiple X-ray co-crystal structures, were synthesized and characterized for concentration-dependent fungal kinase inhibition. Analogs **8**, **9**, and **11** showed improvements in selectivity across the fungal kinome. Most remarkably, phenol **11** was a potent Yck2 inhibitor with improved selectivity over ALK5. MIB/MS competition assays in *C. albicans* revealed **11** as the most selective Yck2/Yck22 fungal kinase inhibitor described to date. Although **LY** and phenol **11** demonstrate whole cell antifungal activity in sensitized backgrounds, their potency appears to be compromised by poor fungal membrane permeability. Additional efforts will be required to improve fungal penetration of these compounds if they are to be developed further for the treatment of *C. albicans* infections.

## Materials and Methods

### General Information

All reagents and solvents used were purchased from commercial sources and were used without further purification. NMR spectra were obtained using Bruker 400 MHz spectrometers at room temperature; chemical shifts are expressed in parts per million (ppm, δ units) and are referenced to the residual protons in the deuterated solvent used. Coupling constants are given in units of hertz (Hz). Splitting patterns describe apparent multiplicities and are designated as s (singlet), d (doublet), t (triplet), q (quartet), m (multiplet), and br s (broad singlet), dd (doublet of doublets), ddd (double double doublet), tt (triplet of triplets). The purity of compounds submitted for biological screening was determined to be ≥95% as measured by HPLC. Analytical thin layer chromatography (TLC) was performed on silica gel plates, 200 μm with an F254 indicator. Column chromatography was performed using RediSep Rf preloaded silica gel cartridges on Isolera one Biotage automated purification systems. Samples for high-resolution mass spectrometry were analyzed with a ThermoFisher Q Exactive HF-X (ThermoFisher, Bremen, Germany) mass spectrometer coupled with a Waters Acquity H-class liquid chromatograph system. Samples were introduced by a heated electrospray source (HESI) at a flow rate of 0.3 mL/min. Electrospray source conditions were set as: spray voltage 3.0 kV, sheath gas (nitrogen) 60 arb, auxillary gas (nitrogen) 20 arb, sweep gas (nitrogen) 0 arb, nebulizer temperature 375 °C, capillary temperature 380 °C, RF funnel 45 V. The mass range was set to 150–2000 m/z. All measurements were recorded at a resolution setting of 120,000. Separations were conducted on a Waters Acquity UPLC BEH C18 column (2.1 × 50 mm, 1.7 *μ*m particle size). LC conditions were set at 95 % water with 0.1% formic acid (A) ramped linearly over 5.0 min to 100% acetonitrile with 0.1% formic acid (B) and held until 6.0 min. At 7.0 min the gradient was switched back to 95% (A) and allowed to re-equilibrate until 9.0 min. Injection volume for all samples was 3 *μ*L. Analytical LC/MS data was obtained using a Waters Acquity Ultrahigh-performance liquid chromatography (UPLC) system equipped with a photodiode array (PDA) detector using the following method: solvent A = Water + 0.2% FA, solvent B = ACN + 0.1% FA, flow rate = 1 mL/min. The gradient started at 95% A for 0.05 min, ramped up to 100% B over 2 min and held for an additional minute at this concentration, before returning to the initial gradient. Compounds were purified by preparative HPLC using an Agilent 1100 equipped with a Phenomenex column (Phenyl-Hexyl, 75 × 30 mm, 5 μm) using the following method: Solvent A: water + 0.05 % TFA; Solvent B: MeOH; flow rate: 70.0 mL/min. LC conditions were set at 90 % (A) ramped linearly over 8.0 min to 100% (B) and held until 10.0 min at 100% B. At 10.0 min the gradient was switched back to 90%. All compounds were 95% pure by NMR analysis.

### Chemistry

#### General Procedure A for Synthesis of LY Analogs

Lepidine (923 µL, 7 mmol, 1.0 equiv) was brought up in anhydrous THF (7.0 mL) to a flame dried flask under argon. The solution was cooled in an ice bath to 0 °C, followed by dropwise addition of 1.5M LiHMDS (14.0 mL, 21 mmol, 3.0 equiv). The mixture was left to stir for 30 minutes. Simultaneously, Methyl picolinate (1.43 mL, 7 mmol, 1.0 equiv) was brought up in anhydrous THF (7.0 mL) to a separate flame dried flask under argon. The methyl picolinate solution was added dropwise via syringe to the reaction mixture over 5 minutes, turning the solution deep green. The reaction was left to run for 18 hours as it warmed to room temperature. Upon completion of reaction, material was rotovaped and diluted in EtOAc (125 mL). The organic layer was washed with NH4Cl (50 mL × 2) and brine (50 mL × 2), dried using Na2SO4, concentrated, and a portion of the crude oil was used in the next step. Ketone intermediate (∼250 mg, 1 mmol, 1.0 equiv) was brought up in anhydrous THF (4.0 mL) and transferred to a flame dried scintillation vial under argon. Hydrazine (1 mmol, 1.0 equiv) was weighed out and transferred to the solution followed by the addition of 3 drops of conc. HCl. The reaction was left to run at 40 °C for 3 hours until full conversion to hydrazide chloride was achieved by TLC/LC-MS. The salt intermediate was filtered, washed with cold EtOAc (20 mL × 2), and placed into a microwave vial. Material was brought up in anhydrous DMF (4.0 mL) and heated in a Biotage microwave at 200 °C for 8 hours. Upon reaction completion, solvent was blown down and material was directly subjected to reversed-phase HPLC. Product appeared as a white solid. Compounds **1** and **2** were synthesized according to General Procedure A.

*4-(5-benzyl-3-(pyridin-2-yl)-1H-pyrazol-4-yl)quinoline* (**1**). Product appeared as a white solid (22.3 mg, 6%). ^1^H NMR (500 MHz, CDCl3) δ 8.82 (d, *J* = 4.7 Hz, 1H), 8.42 (s, 1H), 8.33 (d, *J* = 8.9 Hz, 1H), 7.72 (ddt, *J* = 8.9, 6.9, 1.8 Hz, 1H), 7.57 (dt, *J* = 7.3, 1.4 Hz, 1H), 7.39 – 7.35 (m, 1H), 7.22 (d, *J* = 4.7 Hz, 1H), 7.04 (ddd, *J* = 7.6, 4.8, 1.1 Hz, 1H), 7.02 – 6.94 (m, 4H), 6.86 – 6.78 (m, 2H), 3.89 (d, *J* = 15.6 Hz, 1H), 3.71 (d, *J* = 15.5 Hz, 1H); ^13^C NMR (126 MHz, CDCl3) δ 149.5, 149.2, 138.2, 136.9, 136.5, 131.2, 128.6, 128.3, 128.2, 128.1, 127.9, 127.0, 126.4, 126.1, 125.9, 123.7, 123.2, 123.2, 122.9, 122.2, 120.4, 113.3, 33.0, 31.49. *4-(5-(4-fluorobenzyl)-3-(pyridin-2-yl)-1H-pyrazol-4-yl)quinoline* (**2**). Product appeared as a white solid (8.6 mg, 2%). ^1^H NMR (500 MHz, DMSO) δ 8.87 (d, *J* = 4.3 Hz, 1H), 8.27 (bs, 1H), 8.03 (dd, *J* = 8.5, 1.2 Hz, 1H), 7.68 (ddd, *J* = 8.3, 6.7, 1.5 Hz, 1H), 7.60 (td, *J* = 7.8, 1.8 Hz, 1H), 7.43 (dd, *J* = 8.4, 1.4 Hz, 1H), 7.35 (ddd, *J* = 8.3, 6.8, 1.3 Hz, 1H), 7.30 (d, *J* = 4.3 Hz, 1H), 7.17 – 7.11 (m, 1H), 6.88 (d, *J* = 7.3 Hz, 4H), 3.85 (d, *J* = 15.7 Hz, 1H), 3.69 (d, *J* = 15.7 Hz, 1H); ^13^C NMR (126 MHz, DMSO) δ 162.0, 160.1, 150.5, 149.6, 148.4, 141.3, 137.2, 135.3, 130.4, 130.4, 129.8, 129.6, 127.9, 126.8, 126.1, 123.6, 123.0, 121.0, 115.3, 115.1, 113.9, 31.0.

*General Procedure B for Synthesis of LY Analogs*. Lepidine (923 µL, 7 mmol, 1.0 equiv) was brought up in anhydrous THF (7.0 mL) to a flame dried flask under argon. The solution was cooled in an ice bath to 0 °C, followed by dropwise addition of 1.5M LiHMDS (14.0 mL, 21 mmol, 3.0 equiv). The mixture was left to stir for 30 minutes. Simultaneously, Methyl picolinate (1.43 mL, 7 mmol, 1.0 equiv) was brought up in anhydrous THF (7.0 mL) to a separate flame dried flask under argon. The methyl picolinate solution was added dropwise via syringe to the reaction mixture over 5 minutes, turning the solution deep green. The reaction was left to run for 18 hours as it warmed to room temperature. Upon completion of reaction, material was rotovaped and diluted in EtOAc (125 mL). The organic layer was washed with NH4Cl (50 mL × 2) and brine (50 mL × 2), dried using Na2SO4, concentrated, and the crude oil was used in the next step. Ketone intermediate was brought up in anhydrous THF (14.0 mL) in a flame-dried flask under argon. DMFDMA (4.64 mL, 35 mmol, 5.0 equiv) was added dropwise to the reaction. The reaction was left to stir at 60 °C for 24 hours. Upon completion of reaction, solvent was removed via rotovaping to leave a thick, dark black tar. The crude oil was brought up in 95% ethanol (14.0 mL) and introduced to 60% aq. hydrazine hydrate (7.4 mL, 84 mmol, 12 equiv). The mixture was left to stir overnight at room temperature. Solvent was removed via rotovaping and material was subjected to normal phase flash column chromatography (92% EtOAc/Hex). Product appeared as a gray/white solid (517 mg, 27%). **LY** used in biological experiments/assays was commercially sourced. Otherwise, **LY** intermediate was synthesized in this manner and used crude in later derivatization steps. Compounds **3**, **4**, **7**, **8**, **9**, **10**, **11**, and **12** were synthesized according to General Procedure A.

*4-(1-methyl-3-(pyridin-2-yl)-1H-pyrazol-4-yl)quinoline* (**3**). **LY** intermediate (50 mg, 0.2 mmol, 1.0 equiv) was brought up in anhydrous DMF (0.5 mL) in a flame-dried scintillation vial under argon. 60% sodium hydride in mineral oil (44 mg, 1.1 mmol, 6.0 equiv) was added to the reaction followed by the addition of a few drops of methyl iodide (11 µL, 0.2 mmol, 1.0 equiv). The reaction was left to stir at room temperature for one hour. Upon completion of reaction, solvent was blown down overnight and material was subjected to normal phase flash column chromatography (60% EtOAc/Hex). Product appeared as a clear oil (19.7 mg, 37%). ^1^H NMR (500 MHz, DMSO) δ 8.75 (dd, *J* = 6.7, 4.4 Hz, 1H), 8.03 – 7.97 (m, 2H), 7.95 (dt, *J* = 8.4, 1.6 Hz, 1H), 7.76 – 7.72 (m, 1H), 7.70 – 7.58 (m, 3H), 7.31 (ddd, *J* = 8.2, 6.8, 1.3 Hz, 1H), 7.27 – 7.22 (m, 1H), 7.06 (ddd, *J* = 7.4, 4.8, 1.3 Hz, 1H), 3.96 (s, 3H); ^13^C NMR (126 MHz, DMSO) δ 152.5, 150.3, 149.2, 148.5, 148.4, 141.1, 137.1, 133.9, 129.7, 129.4, 127.6, 126.5, 126.5, 122.8, 122.8, 121.8, 116.5, 39.5.

*4-(1-benzyl-3-(pyridin-2-yl)-1H-pyrazol-4-yl)quinoline* (**4**). **LY** intermediate (65 mg, 0.2 mmol, 1.0 equiv) was brought up in anhydrous DMF (0.5 mL) in a flame-dried scintillation vial under argon. 60% sodium hydride in mineral oil (57 mg, 1.4 mmol, 6.0 equiv) was added to the reaction followed by the dropwise addition of benzyl bromide (28 µL, 0.2 mmol, 1.0 equiv). The reaction was left to stir at room temperature for one hour. Upon completion of reaction, solvent was blown down overnight and material was subjected to normal phase flash column chromatography (85% EtOAc/Hex). Product appeared as white crystals (50.2 mg, 58%). ^1^H NMR (500 MHz, DMSO) δ 9.17 (d, *J* = 5.4 Hz, 1H), 8.50 (s, 1H), 8.25 (dd, *J* = 8.7, 1.0 Hz, 1H), 8.09 (dt, *J* = 4.7, 1.4 Hz, 1H), 8.02 (ddd, *J* = 8.4, 6.9, 1.3 Hz, 1H), 7.97 (dd, *J* = 8.6, 1.3 Hz, 1H), 7.90 (dt, *J* = 8.0, 1.2 Hz, 1H), 7.87 – 7.82 (m, 2H), 7.69 (ddd, *J* = 8.4, 6.9, 1.2 Hz, 1H), 7.49 – 7.39 (m, 4H), 7.39 – 7.33 (m, 1H), 7.23 (ddd, *J* = 7.5, 4.9, 1.3 Hz, 1H), 5.58 (s, 2H); ^13^C NMR (126 MHz, DMSO) δ 151.1, 149.7, 148.8, 148.7, 145.6, 140.4, 138.1, 137.1, 134.8, 133.6, 129.2, 129.2, 128.5, 128.4, 127.6, 127.5, 123.6, 123.3, 123.2, 122.2, 119.6, 117.3, 115.5, 115.0, 112.7, 56.0. *2-(4-(pyridin-4-yl)-1H-pyrazol-3-yl)pyridine* (**7**). Product appeared as a white solid (8.2 mg, 1%). ^1^H NMR (500 MHz, DMSO) δ 8.57 (d, *J* = 4.8 Hz, 1H), 8.49 – 8.43 (m, 2H), 8.14 (s, 1H), 7.88 (td, *J* = 7.7, 1.8 Hz, 1H), 7.69 (s, 1H), 7.43 – 7.35 (m, 3H); ^13^C NMR (126 MHz, DMSO) δ 149.8, 149.6, 141.6, 137.4, 123.6, 123.3, 123.2, 118.0.

*4-(3-(pyridin-2-yl)-1H-pyrazol-4-yl)pyrimidine* (**8**). Product appeared as a white solid (15.5 mg, 2%). ^1^H NMR (500 MHz, DMSO) δ 9.06 (d, *J* = 1.4 Hz, 1H), 8.68 – 8.58 (m, 2H), 8.34 (s, 1H), 7.91 (td, *J* = 7.7, 1.8 Hz, 1H), 7.86 – 7.80 (m, 1H), 7.65 (dd, *J* = 5.4, 1.5 Hz, 1H), 7.43 (ddd, *J* = 7.5, 4.8, 1.3 Hz, 1H); ^13^C NMR (126 MHz, DMSO) δ 159.8, 158.8, 157.1, 149.5, 137.4, 123.9, 119.5, 118.4.

*4-(3-(5-fluoropyridin-2-yl)-1H-pyrazol-4-yl)quinoline* (**9**). Product appeared as a grey solid (10.1 mg, 3%). ^1^H NMR (500 MHz, DMSO) δ 8.87 (d, *J* = 4.4 Hz, 1H), 8.06 (dd, *J* = 8.8, 1.3 Hz, 1H), 7.77 – 7.70 (m, 2H), 7.47 (ddd, *J* = 8.2, 6.9, 1.3 Hz, 1H), 7.36 (d, *J* = 4.4 Hz, 1H), 7.33 – 7.27 (m, 2H), 7.07 (s, 2H). ^13^C NMR (126 MHz, DMSO) δ 163.0, 161.1, 150.6, 148.7, 140.7, 129.9, 129.5, 129.4, 127.3, 127.1, 126.2, 123.2, 116.0, 115.9, 114.6.

*4-(3-(5-methylpyridin-2-yl)-1H-pyrazol-4-yl)quinoline* (**10**). Product appeared as a grey solid (6.9 mg, 4%). ^1^H (500 MHz, DMSO) δ 9.16 (d, *J* = 5.4 Hz, 1H), 8.25 (d, *J* = 8.4 Hz, 2H), 8.13 (s, 1H), 8.02 (t, *J* = 7.5 Hz, 1H), 7.98 (d, *J* = 8.5 Hz, 1H), 7.81 (d, *J* = 5.3 Hz, 1H), 7.75 – 7.67 (m, 2H), 7.64 (d, *J* = 8.3 Hz, 1H), 2.24 (s, 3H); ^13^C NMR (126 MHz, DMSO) δ 149.5, 148.6, 147.2, 145.8, 141.0, 139.2, 133.5, 129.2, 127.5, 123.5, 122.1, 119.6, 117.3, 115.0, 114.7, 112.7, 18.1.

*2-(4-(quinolin-4-yl)-1H-pyrazol-3-yl)phenol* (**11**). 2-(4-(quinolin-4-yl)-1H-pyrazol-3-yl)phenol (250 mg, 0.6 mmol, 1.0 equiv) was prepared under General Procedure B conditions. This PMB-protected intermediate was brought up in DCM (1.2 mL) in a scintillation vial. TFA (473 µL, 6.1 mmol, 10.0 equiv) was added to the mixture, and the reaction was left to stir at room temperature for 3 hours. Upon completion of reaction, solvent was blown down and material was directly loaded for reversed-phase prep-HPLC (1% NH4OH, Rf = 4.2 min). Product appeared as a white crystal (26.9 mg, 15%). ^1^H NMR (500 MHz, DMSO) δ 10.31 (s, 0.5H), 9.73 (s, 0.5H), 8.81 (d, *J* = 41.7 Hz, 1H), 8.18 (s, 0.5H), 8.09 – 7.97 (m, 1H), 7.91 (d, *J* = 8.5 Hz, 0.5H), 7.82 – 7.67 (m, 1.5H), 7.47 (s, 1H), 7.28 (d, *J* = 61.2 Hz, 1H), 7.09 (d, *J* = 29.8 Hz, 1H), 7.01 – 6.86 (m, 1H), 6.80 (d, *J* = 8.5 Hz, 1H), 6.67 (s, 1H), 6.54 (s, 0.5H).^13^C NMR (126 MHz, DMSO) δ 150.53, 148.59, 141.34, 140.28, 131.20, 130.85, 130.36, 129.78, 129.40, 128.95, 127.03, 126.64, 126.36, 122.60, 122.02, 119.36, 119.13, 116.65, 116.29.

*4-(3-(furan-2-yl)-1H-pyrazol-4-yl)quinoline* (**12**). Product appeared as a white solid (24.0 mg, 3%). ^1^H NMR (500 MHz, DMSO) δ 8.84 (d, *J* = 4.3 Hz, 1H), 8.01 (d, *J* = 8.4 Hz, 1H), 7.92 (bs, 1H), 7.67 (dd, *J* = 17.6, 8.3 Hz, 2H), 7.43 (t, *J* = 7.6 Hz, 2H), 7.36 (d, *J* = 4.4 Hz, 1H), 6.32 (s, 1H), 6.13 (s, 1H); ^13^C NMR (126 MHz, DMSO) δ 150.5, 148.5, 143.1, 140.2, 129.9, 129.8, 127.5, 127.1, 126.1, 123.1, 114.2, 111.8, 107.8

*4-(2-(pyridin-2-yl)-1H-pyrrol-3-yl)quinoline* (**5**). 1-(pyridin-2-yl)-2-(quinolin-4-yl)ethan-1-one (350 mg, 1.4 mmol, 1.0 equiv) was brought up in dry DMSO (1.4 mL) in a flame-dried 5.0 microwave vial under argon. The vessel was placed into an cool water bath. 1.0M LiHMDS in THF (1.6 mL, 1.5 mmol, 1.1 equiv) was added dropwise to the mixture and left to stir for 30 minutes. At this point, allyl bromide (171 µL, 2.0 mmol, 1.4 equiv) was added dropwise in 2 increments over 30 minutes separated by 20 minutes between additions. The reaction was left to stir for 3 hours, although some starting material did remain. Material was diluted in EtOAc (20 mL) and the organic layer was washed with NH4Cl (15 mL × 2) and brine (15 mL × 2), dried using Na2SO4, concentrated, and subjected to normal phase chromatography (30% EtOAc/Hex). 1-(pyridin-2-yl)-2-(quinolin-4-yl)pent-4-en-1-one appeared as a clear oil (95 mg, 23%). ^1^H NMR (500 MHz, CDCl3) δ 8.72 (dd, *J* = 4.7, 1.4 Hz, 1H), 8.47 – 8.42 (m, 1H), 8.41 (d, *J* = 8.3 Hz, 1H), 8.03 (dd, *J* = 8.5, 1.4 Hz, 1H), 7.94 (dt, *J* = 7.9, 1.2 Hz, 1H), 7.66 (td, *J* = 7.7, 1.8 Hz, 1H), 7.62 (ddd, *J* = 8.3, 6.8, 1.4 Hz, 1H), 7.53 (ddd, *J* = 8.4, 6.8, 1.4 Hz, 1H), 7.27 (ddt, *J* = 9.3, 4.9, 3.3 Hz, 2H), 5.75 – 5.64 (m, 1H), 5.00 (dq, *J* = 17.0, 1.6 Hz, 1H), 4.85 (dt, *J* = 9.9, 1.4 Hz, 1H), 3.02 – 2.94 (m, 1H), 2.67 – 2.58 (m, 1H); ^13^C NMR (126 MHz, CDCl3) δ 200.1, 152.4, 149.8, 149.0, 148.4, 145.8, 137.0, 135.2, 130.0, 129.3, 127.4, 127.2, 126.7, 124.06, 122.63, 120.14, 117.14, 45.25, 37.06.

1-(pyridin-2-yl)-2-(quinolin-4-yl)pent-4-en-1-one (95 mg, 0.3 mmol, 1.0 equiv) was brought up in MeOH (1.0 mL) and DCM (0.1 mL) in a scintillation vial. The reaction was cooled to -78 C. Using an ozonylizer, the reaction was sparged with ozone for 2 minutes, turning the solution bright blue. At this point, the reaction vessel was sparged with argon for 20 minutes and a vent needle to remove excess ozone. Dimethyl sulfide (73 µL, 1.0 mmol, 3.0 equiv) was added dropwise, and the reaction was left to stir overnight and coom to room temperature. Solvents were removed via rotovaping, and the material was used in the next step without further purification. This material was separated into batches for synthesis of **5** and **6**. The diketone intermediate (50.0 mg, 0.2 mmol, 1.0 equiv) was brought up in glacial acetic acid (1.2 mL) in a 5.0 mL microwave vial alongside ammonium acetate (226 mg, 2.9 mmol, 17 equiv). The resulting mixture was heated using the Biotage microwave at 120 °C for 2 hours. Material was diluted in EtOAc (15 mL) and aq. NaHCO3 (15 mL). Solid Na2CO3 was added until effervescence ceased. The organic phase was washed with NaHCO3 (10 mL) and brine (20 mL), dried using Na2SO4, concentrated, and material was directly loaded for reversed-phase prep-HPLC (1% NH4OH, Rf = 3.9 min). Product appeared as a white solid (9.5 mg, 20% overall). ^1^H NMR (500 MHz, DMSO) δ 9.10 (d, *J* = 5.3 Hz, 1H), 8.39 (dt, *J* = 4.7, 1.5 Hz, 1H), 8.21 (dd, *J* = 8.6, 1.1 Hz, 1H), 8.04 (dd, *J* = 8.6, 1.3 Hz, 1H), 7.98 (ddd, *J* = 8.4, 6.9, 1.4 Hz, 1H), 7.77 (d, *J* = 5.3 Hz, 1H), 7.67 (ddd, *J* = 8.4, 6.9, 1.2 Hz, 1H), 7.54 (td, *J* = 7.8, 1.8 Hz, 1H), 7.20 (t, *J* = 2.7 Hz, 1H), 7.13 (ddd, *J* = 7.4, 4.8, 1.1 Hz, 1H), 7.07 (d, *J* = 8.0 Hz, 1H), 6.46 (t, *J* = 2.6 Hz, 1H); ^13^C NMR (126 MHz, DMSO) δ 158.8, 158.5, 150.1, 149.6, 146.2, 137.3, 133.1, 130.2, 128.8, 127.8, 127.5, 124.2, 123.0, 122.0, 121.8, 120.9, 118.1, 113.3.

*4-(2-(pyridin-2-yl)furan-3-yl)quinoline* (**6**). The diketone intermediate (50.0 mg, 0.2 mmol, 1.0 equiv) was brought up in glacial acetic acid (0.3 mL) in a 5.0 mL microwave vial conc. HCl (0.8 mL). The resulting mixture was heated using the Biotage microwave at 120 °C for 2 hours. Following heating, the vial was placed into an ice bath, and a saturated solution of aq. K2CO3 was added until it became basic by litmus paper. The aqueous phase was extracted with EtOAc (10 mL × 4), dried using Na2SO4, concentrated, and material was directly loaded for reversed-phase prep-HPLC (1% TFA, Rf = 2.5 min). Product appeared as a white solid (8.5 mg, 18% overall). ^1^H NMR (500 MHz, DMSO) δ 9.05 (d, *J* = 4.7 Hz, 1H), 8.18 – 8.15 (m, 2H), 8.12 (d, *J* = 1.8 Hz, 1H), 7.87 (ddd, *J* = 8.3, 6.8, 1.4 Hz, 1H), 7.80 (td, *J* = 7.7, 1.7 Hz, 2H), 7.68 (d, *J* = 4.7 Hz, 1H), 7.64 (dt, *J* = 8.0, 1.1 Hz, 1H), 7.59 (ddd, *J* = 8.3, 6.8, 1.2 Hz, 1H), 7.19 (ddd, *J* = 7.6, 4.8, 1.1 Hz, 1H), 6.91 (d, *J* = 1.8 Hz, 1H); ^13^C NMR (126 MHz, DMSO) δ 149.68, 149.31, 148.8, 148.8, 144.64, 137.61, 131.32, 127.99, 127.52, 127.17, 126.58, 123.23, 122.99, 120.51, 120.29, 116.00.

### X-ray Crystallography

Yck2 for crystallography was cloned and purified as previously reported^3^. Crystallization was performed at RT using the sitting drop method. Crystals of *apo*Yck2 were grown with 1 μL protein at 14 mg/mL and 1 μL reservoir solution 0.1 M Tris pH 8, 25 mM magnesium chloride and 20% (w/v) PEG3350. 1 μL of LY dissolved in DMSO were soaked into *apo* Yck2 crystals and incubated for 24 hours. Crystals were cryoprotected in Paratone oil. X-ray diffraction data at 100 K was collected at a Rigaku MicroMax-003 with an Eiger R 1 M detector. Diffraction data was reduced using CrysAlisPro^24^ and the *apo* Yck2 structure (PDB 6U69) was used for Molecular Replacement (MR) to obtain initial phases using Phenix.phaser.^19^ Refinement was completed with Phenix.refine and Coot.^20^ The presence of compounds was readily apparent in Fo − Fc maps after MR. All B-factors were refined, and TLS parameterization was included in final rounds of refinement. Geometry was verified using Phenix.molprobity and the wwPDB validation server, and structures were deposited to the Protein Databank with accession number PDB: 9NZK. X-ray crystallographic statistics are provided in SI Table 2.

### Biology

#### HEK293 Cell Lysate

Expi293 cells (ThermoFisher Scientific were grown to approximately 3.0×10^6^ cells/mL (log phase growth, well below saturation) in Expi293 medium following standard protocols from the manufacturer. The cells were collected by centrifugation at 300xg at 4 °C for 20 minutes. The pellets were gently washed thrice with ice cold PBS. Then the pellets were resuspended in cold PBS and portioned into 100–150×10^6^ cell aliquots. Each aliquot was pelleted one last time, the PBS siphoned off, and the pellet frozen at -80 °C until processing. Each aliquot of 100–150×10^6^ cells gave roughly 5 mg total protein in processing. Cell pellets were thawed on ice and introduced to fresh MIB lysis buffer (50 mM HEPES, 150 mM NaCl, 0.5% Triton X-100, 1mM EDTA, 1mM EGTA, 10 mM NaF, 2.5 mM NaVO4, pH 7.5, Phosphatase Inhibitor Cocktail 2 & 3 [Sigma P5726 and P0044], cOmplete Protease Inhibitor Cocktail [EDTA free, Roche). Cells were incubated in MIB lysis buffer for 10 min on ice with tube inverting every 2 min. Cells were sonicated using a Qsonica ultrasonicator at 125 W, 20 kHz, 35% amplitude, in 3×10 sec treatments, allowing sample to cool on ice for 30 sec between pulses. Cell lysate was clarified through 0.20 mm syringe filters (Corning, 431219) into a single 15 mL prechilled Falcon tube. Protein concentration was determined by Bradford method.

#### Fungal Strain and Culture Conditions

Archives of all strains were maintained at -80 °C in 25% glycerol. Strains were grown in standard conditions at 30 °C in YPD (1% yeast extract, 2% peptone, 2% dextrose), unless otherwise indicated. Active cultures were maintained on solid YPD (2% agar) at room temperature no longer than 5 days. Fungal strains include: CaLC2742^25^, CaLC249^25^, CaSS1 *tetO-YCK2/yck2Δ*^26^, and CaSS1 *tetO-ERG6/erg6Δ*.^27^

#### C. albicans Cell Lysate

Archives of all strains were maintained at -80 °C in 25% glycerol. Strains were grown in standard conditions at 30 °C in YPD (1% yeast extract, 2% peptone, 2% dextrose), unless otherwise indicated in RPMI (10.4 g/L RPMI powder with L-glutamine (Gibco), 165 mM MOPS, 2% glucose, 5 mg/mL histidine, pH 7), or SD (2% glucose, 6.7 g/L yeast nitrogen base without amino acids). *C. albicans* strain SC5315 was grown to mid-late log phase (OD 3–4) in four liters YPD medium at 30 °C and harvested by centrifugation at 3,000 g. The pellet was transferred into a 60 mL syringe and squeezed into liquid nitrogen. Cells were disrupted by grinding in liquid nitrogen using mortar-pestle. Each gram of the resulting powder was dissolved in 2 mL of 1.5x MIB buffer (75 mM HEPES-NaOH pH 7.5, 225 mM NaCl, 0.75% Triton X-100, 1.5 mM EDTA, 1.5 mM EGTA, 15 mM NaF, 3.75 mM Na3VO4, 7.5%(v/v) glycerol) supplemented with cOmplete proteinase inhibitor (Roche; 2x of the recommended concentration) and phosphatase inhibitor cocktail II (Sigma). The lysate was sonicated for 20 min (10 sec on; 10 sec off) at 30% amplitude using a Misonix S-4000 dual horn sonicator with 3/4-inch probes and spun at 17,000 rpm in a JA25.50 rotor for 10 min. The supernatant was further clarified by a 90 min centrifugation in a Ti45 rotor at 40,000 rpm. Protein concentration was determined by Bradford method using a 1:10 diluted lysate in 1x TE buffer (10 mM Tris-HCl pH 7.5, 1 mM EDTA).

#### Biochemical Assay

Kinase assays were performed using the ADP-Glo kinase assay kit (Promega) in solid white 384 well plates (Corning). Assays with were performed in kinase buffer (1x: 2 mM NaHEPES pH 7.5, 650 mM KCl, 50 mM MgCl2, 25 mM b-glycerophosphate) supplemented with 2 µg casein kinase 1 peptide substrate (SignalChem) per reaction, 0.05 µg pure p38a (Promega V2701), ∼0.115 μg purified recombinant Yck2 kinase domain, and 0.05 µg purified CK1a (Abcam) per reaction as indicated. Each kinase inhibitor of interest was added in a two-fold dilution series at the concentrations indicated, followed by the addition of ATP at 20 µM (Yck2 KmATP), 10 µM (CK1a) or 80 µM (p38a), respectively. Assays were performed in 10 μL reactions (*n* = 3) and incubated for 30 min at 30 °C. ADP-Glo kinase assay reagents (Promega) were applied to assay wells per manufacturer’s instructions. Luminescence was measured with a TECAN Spark multimode microplate reader, and background luminescence was subtracted from reaction wells from control wells incubated with all reaction reagents except for the relevant kinase enzymes. IC50 values were calculated, and data plotted using GraphPad Prism 9’s Nonlinear fit function.

#### TGFBR1 Kinase Assay

TGFBR1 enzyme inhibition was determined using the Eurofins KinaseProfiler Radiometric Assay at the Km ATP (Cat#: 14-912KP). TGFBR1 (h) is incubated with 8 mM MOPS pH 7.0, 0.2 mM EDTA, 1 mM MnCl2, 2 mg/mL casein, 10 mM MgAcetate and [gamma-33P]-ATP (specific activity and concentration as required). The reaction is initiated by the addition of the Mg/ATP mix. After incubation for 40 minutes at room temperature, the reaction is stopped by the addition of phosphoric acid to a concentration of 0.5%. An aliquot of the reaction is then spotted onto a filter and washed four times for 4 minutes in 0.425% phosphoric acid and once in methanol prior to drying and scintillation counting.

#### Anti-fungal Assay

All tested compounds were dissolved in DMSO. Drug susceptibility assays were performed in 384-well plates in a final volume of 0.04 mL/well (∼1 × 10^3^ cells/mL *C. albicans*) with two-fold dilutions of each compound in YPD or RPMI medium (10.4 g/L RPMI powder with L-glutamine (Gibco), 165 mM MOPS, 2% glucose, 5 mg/mL histidine, pH 7), as indicated. Plates were incubated in the dark at 30 °C or 37 °C with 5% CO2 under static conditions. Absorbance was measured at 600 nm after the indicated incubation times using a spectrophotometer (Molecular Devices) to assess growth. Growth was normalized to the no-drug controls. All assays represent average values of technical duplicates, and were performed with two biological replicates to confirm reproducibility. Data was quantitatively displayed as heat maps using the program Java TreeView3. For assays utilizing the conditional expression strains from the Gene Replacement and Conditional Expression (GRACE) collection, strains were grown overnight in the presence and absence of the indicated doxycycline concentration(s) to repress gene expression.

### Chemoproteomics

#### MIB/MS Assay

Protein lysate containing 5 mg total protein in 4.0 mL MIB lysis buffer was mixed with 4 µL DMSO or test compound in DMSO followed by 1 h incubation on ice. Bio-Rad Poly-Prep Columns (Product # 731-1550) were prepared for each sample with 350 µL of a 50% slurry of a total MIB matrix mix; kinobead matrix consisting of Shokat, PP58, Purvalanol B, UNC-21474, VI-16832, and Ctx-0294885. Each bead type used in the MIB/MS assay consisted of an inhibitor conjugated to Sepharose 4B matrix support in 20% aq. ethanol. Column equilibration was performed with high salt MIB wash buffer (50 mM HPES, 1M NaCl, 0.5% Triton X-100, 1 mM EDTA, 1 mM EGTA, pH 7.5). Cell lysate samples were brought to 1M NaCl. Lysate samples were pipetted onto columns, allowing flow through to pass. Columns were flushed with high salt MIB wash buffer, low salt MIB wash buffer (50 mM HPES, 150 mM NaCl, 0.5% Triton X-100, 1 mM EDTA, 1 mM EGTA, pH 7.5), and 500 µL 0.1% SDS in low salt MIB wash buffer. Kinase elution was performed twice by adding 500 µL MIB elution buffer (0.5% w/v SDS, 0.1 M Tris-HCl, pH 7.25) followed by heating at 95 °C for 10 min. The eluted proteins were reduced with 5mM DL-Dithiothreitol followed by cysteine alkylation using 18 mM iodoacetamide in the dark. The alkylation reaction was quenched by adding DL-Dithiothreitol for a final concentration of 10 mM. Protein concentration was performed with Amicon Millipore Ultra-4 10K cutoff spin columns (Cat# UFC801008) followed by methanol/chloroform extraction. The dry pellets were reconstituted within100 μL of 50 mM HEPES, pH 8.0 buffer. Trypsin digestion, using 1.2 μg sequencing-grade Modified Trypsin (Promoega V5111), was performed overnight at 37 °C.Ethyl acetate extraction was performed on the tryptic peptides followed by sample clean up using C-18 Peptide Desalting Spin Columns (Pierce, Cat# 89852). All samples were made n=1 and analyzed by LC-MS in technical triplicate.

#### LC-MS/MS Analysis

All samples were subjected to LC-MS/MS analysis using a VanquishNeo coupled to an Orbitrap Astral mass spectrometer (Thermo Scientific). Samples were injected onto an Ionoptics Aurora series 3 C18 column (75 μm id × 25 cm, 1.7 μm particle size; Ionopticks) and separated over a 30 min method. The gradient for separation consisted of 5–45% mobile phase B at a 300 nl/min flow rate, where mobile phase A was 0.1% formic acid in water and mobile phase B consisted of 0.1% formic acid in 80% ACN. Astral was operated in product ion scan mode for Data Independent Acquisition (DIA). MS scan range was m/z 380–980; the resolution was set to 240,000 with a 5 ms maximum injection time and AGC target of 500%. Following the full MS scan, the DIA scans were collected with AGC target set to 500%; maximum injection time set to 3 ms; precursor mass range set to 380–980 m/z; the isolation window was set to 3 m/z. Raw data files were processed using Spectronaut (version 19; Biognosys) and searched against the *C. albicans* strain SC5314 database (UP000000559, containing 6,039 entries, downloaded May 2024) appended with a common contaminants database (245 sequences) using the Pulsar search algorithm. Enzyme specificity was set to trypsin, up to two missed cleavage sites were allowed; methionine oxidation, and Protein N-term acetylation were set as variable modifications and carbamidomethylation of Cys was set as a fixed modification. Data was normalized using the global normalization strategy and no imputation was performed. All data were filtered at a 1% false discovery rate (FDR), and proteins were filtered for a minimum of 1 peptide. Log2 transformation of the protein quantities was performed. Log2 fold change (FC) ratios were calculated using the Log2 LFQ intensities of drug treated sample compared to control.

## Supporting information

Supplemental Information 1 - MIB/MS Data

## Supporting Information

Table S1, Sequence alignment of represented kinases; Table S2, X-ray crystallographic statistics for co structures between *C. albicans* Yck2 and **LY**; Figure S1, fungal protein kinome (103) and their representative kinase families based on Goswami and authors; Figure SX–Y, Compound characterization; File S1 MS proteomics data.

## Author Contributions

The manuscript was written through contributions of all authors.

## Acknowledgments

We would like to acknowledge Aurora Cabrera, Laura E. Herring, and Lee M. Graves of the UNC Lineberger Comprehensive Cancer Center, Department of Pharmacology, and UNC Metabolomics and Proteomics core for their facilities, reagents, and aid in conducting the MIB/MS competition experiments as well as manuscript drafting. We’d also like to thank John Forsberg and Nathaniel I. Nicely of the UNC Protein Expression and Purification Core contributed HEK293 cell pellets for the MIB/MS experiments. We thank Merck and Genome Canada for making the original *Candida albicans* GRACE mutant collections available.

