## Supporting Information for "Structure-activity Relationship for Diarylpyrazoles as Inhibitors of the Fungal Kinase Yck2"

### Table of Contents

| | Page |
|---|---|
| Table S1 | S2 |
| Table S2 | S3 |
| Figure S1 | S4 |
| Compound Characterization | S5-X |

**Table S1.** Sequence alignment of represented kinases. Key structural elements are highlighted in yellow. P-loop, glycine rich flexible loop; CatLys, catalytic lysine; α-C, αC helix; Gate-Hinge, gatekeeper residue and hinge binding motif; DFG, phosphate binding DFG motif.

| | | |
|---|---|---|
| | **P-loop** (GKGRFG…) **CatLys** (VAVK…) | |
| ALK5 HUMAN | IYDM--TTSGSGSGLPLLVQRTIARTIVLQESIGKGRFGEVWRGK--WRGEEVAVKIFSS | 236 |
| YCK2 CALBI | MNHSTSSSNGNGSNNSS----VVGLHYKIGKKIGEGSFGVIFEGTNIINGVPVAIKFEPR | 77 |
| CK1A HUMAN | ------MASSSGSKAEF----IVGGKYKLVRKIGSGSFGDIYLAINITNGEEVAVKLESQ | 50 |
| MK14 HUMAN | SQERPTFYRQELNKTIW----EVPERYQNLSPVGSGAYGSVCAAFDTKTGLRVAVKKLSR | 57 |
| HOG1 CALBI | MSADGEFTRTQIFGTVF----EITNRYTELNPVGMGAFGLVCSAVDRLTGQNVAVKKVMK | 56 |
| Consensus | ` . : :* * :* : . * **:*` |  |
| | α-C (EAEIY…) Gate-Hinge (WLVSDYHE…) | |
| ALK5 HUMAN | REERSW-----FREAEIYQTVMLRHENILGFIAADNK--DNGTWTQLWLVSDYHE--HGS | 287 |
| YCK2 CALBI | KTEAP----QLRDEYRTYK--HLQGCDGIPNAYYFGQ-----EGLHNILVIDLLGPSLED | 126 |
| CK1A HUMAN | KARHP----QLLYESKLYK--ILQGGVGIPHIRWYGQ-----EKDYNVLVMDLLGPSLED | 99 |
| MK14 HUMAN | PFQSIIHAKRTYRELRLLK--HMKHENVIGLLDVFTPARSLEEFNDVYLVTHLMGADLNN | 115 |
| HOG1 CALBI | PFSTSVLAKRTYRELKLLK--HLKHENLITLDDIFIS-----PLEDIYFVNELQGTDLHR | 109 |
| Consensus | ` * . : :: : :* .` |  |
| | DFG (DLG/DFG) | |
| ALK5 HUMAN | ---TCCIADLGLAVRHDSATDTIDIAPNHRVGTKRYMAPEVLDDSINMKHF------ESF | 396 |
| YCK2 CALBI | DENNVHLIDFGMAKQYRDPRTK-------QHI--PYREKKSLSGTARYMSINTHLGREQS | 228 |
| CK1A HUMAN | HCNKLFLIDFGLAKKYRDNRTR-------QHI--PYREDKNLTGTARYASINAHLGIEQS | 199 |
| MK14 HUMAN | --CELKILDFGLARHTDDEMTG-------YVATRWYRAPEIML---NWMHY--------N | 201 |
| HOG1 CALBI | --CDLKICDFGLARLQDPQMTG-------YVSTRYYRAPEIML---TWQKY--------D | 195 |
| Consensus | ` : *:*:* * : :` |  |

**Table S2.** X-ray crystallographic statistics for co structures between *C. albicans* Yck2 and **LY**

| **Structure** | **Yck2 + LY** |
|---|---|
| **PDB code** | **9NZK** |
| **Data collection** | |
| Space group | R 3 |
| Cell dimensions | |
| *a*, *b, c* (Å) | 91.53, 91.53, 118.824 |
| α, β, γ, (°) | 90, 90, 120 |
| Resolution (Å) | 29.95 – 1.70 |
| $R_{merge}$[a] | 0.050 (0.622)* |
| $CC_{1/2}$* | 0.999 (0.445) |
| $F / \sigma F$ | 19.25 (0.48) |
| Completeness (%) | 99.8 (98.8) |
| Redundancy | 3.4 (1.7) |
| **Refinement** | |
| Resolution, Å | 29.95 – 1.70 |
| No. reflections: working, test | 40743, 1949 |
| $R_{work}/R_{free}$[c] | 17.6/21.1 (35.5/39.1) |
| No. atoms | |
| Protein | 2469 |
| Inhibitor | 21 |
| Solvent | 2 |
| Water | 357 |
| *B*-factors | |
| Protein | 39.0 |
| Inhibitor | 30.2 |
| Solvent | 47.1 |
| Water | |
| R.m.s. deviations | |
| Bond lengths (Å) | 0.011 |
| Bond angles (°) | 1.112 |

*Values in parentheses are for highest-resolution shell.

**Figure S1.** Fungal protein kinome (103) and their representative kinase families based on Goswami and authors. Green circles indicate kinases covered by MIB/MS (86), with empty circles showing those not covered (17).

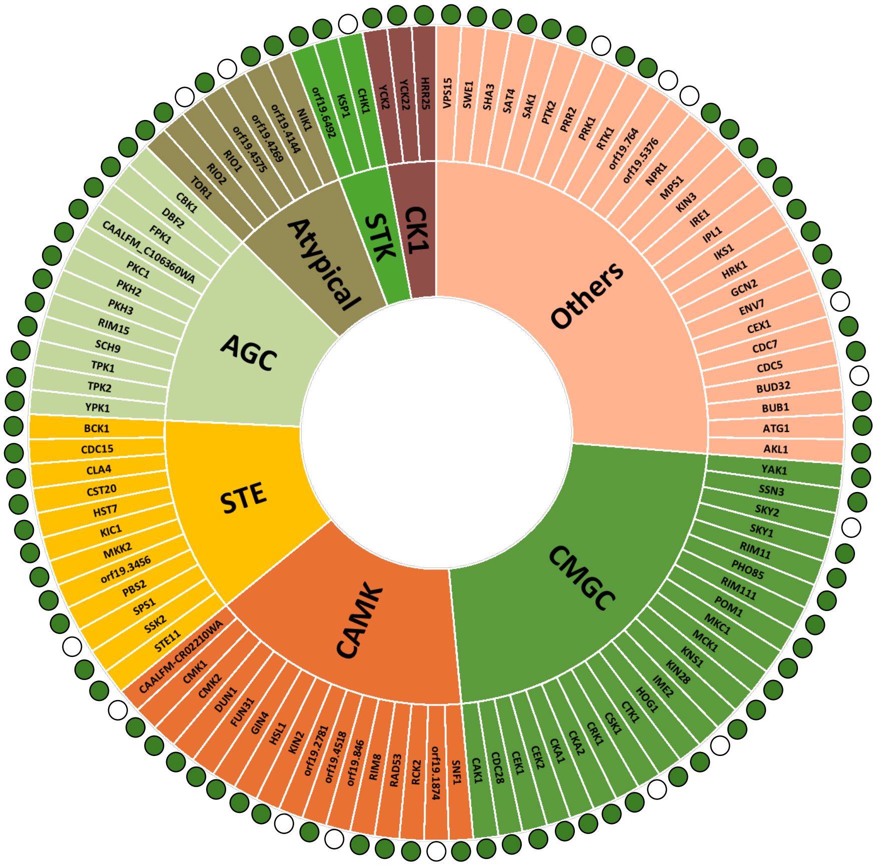
